# Autonomous Spatial Transcriptomics Analysis (ASTA): Demonstrating Performance Improvements through Clustering, Biological Annotation, and AI-Driven Discovery

**DOI:** 10.64898/2026.08.10.743848

**Authors:** Maiqi Zhang, Minjun Roe, Chris Pollett, William B. Andreopoulos

## Abstract

Spatial transcriptomics keeps measurement of gene expression while preserving spatial context, yet traditional analysis methods face challenges in computational efficiency, biological interpretability, and autonomous discovery. This project presents a framework solving these issues through three parts: (1) an ensemble clustering system achieving 66.7% improvement over baseline average and 23.9% over best single method with silhouette score of 0.540 and statistical significance (*p* = 0.0032, Cohen’s *d* = 1.82); (2) a knowledge-based clustering framework that annotates 88.6% of cells across 8 ovarian cell types using 428 marker genes; and (3) a GPT-4o-mini-powered autonomous agent that generated 3 biological hypotheses with validations.

## 1 Introduction to Spatial Transcriptomics, Background and Motivation

Gene expression analysis has been important for molecular biology research [1]. All of the cells within a multicellular organism contain the same genome, but cells show different neurons transmit electrical signals. This complete set of genes expressed in a cell is called a transcriptome [1]. It is a molecular fingerprint that defines cellular identity and function.

Transcriptomic methods have been driven by the rapid development of new technologies with an improved sensitivity and economy [2]. However, traditional methods share a similar limitation, which is that they require tissue dissociation, which eliminates the spatial information of cells within the tissue. This loss of positional information is significant because cellular behavior is easily influenced by spatial context.

Spatial transcriptomics solves this issue by keeping both gene expression and gene spatial coordinates within the tissue [3]. This new technology allows researchers to see not just which genes are active, but also exactly where that activity happens inside the tissue. [4].

### 1.1 Current Technological Landscape

Since its recognition in recent years, spatial transcriptomics has been widely used in different research applications [4]. For example, it can be used to map the structure and development of the same organ in different evolutionary species, providing a deeper understanding of the evolution of organs at the cellular level and a basis for studying new tissues and their functions.

Stereo-seq is one of the techniques in spatial transcriptomics[5]. It allows scientists to visualize and analyze the complexity of tumors at an unprecedented level of detail [5].

However, the analysis of spatial transcriptomics data poses some challenges, such as computational challenges and lack of biological meaning. More specifically, it is facing the following three limitations:

#### 1.1.1 Limitation 1: Single-Method Clustering Approaches

Relying on a single clustering algorithm such as K-means alone or hierarchical clustering alone often fail to capture the complexity of heterogeneous tissue samples, because the tissue performance varies across different types and experimental conditions. A single clustering method that performs well on one dataset does not mean it will perform well on the other dataset. This instability leads to inconsistent results and makes it difficult to compare findings across different researches.

#### 1.1.2 Limitation 2: Lack of Biological Meaning Integration

Computational methods are usually implemented seperately from biological knowledge, which leads to only show mathematically optimal results but lacking biological meaning. This requires manual interpretation and validation by domain experts, which creates a bottleneck in the analysis pipeline and limits the scalability of spatial transcriptomic research.

#### 1.1.3 Limitation 3: Human-Dependent Analysis Workflows

From data preprocessing, research implementation, and result interpretation, human decision-making is critical at each stage, which potentially leads to personal bias, unintentional error, and random variability. This dependence on human expertise also creates a bottleneck in spatial transcriptomic studies.

### 1.2 Project Objectives

This project solves the issues described above through three components: (1) an ensemble clustering system that combines K-means, hierarchical clustering, and Gaussian mixture models through a weighted co-association matrix to produce more robust and stable cluster assignments than any single method alone; (2) a knowledge-based biological annotation framework that integrates ovarian-specific marker genes and biological pathways to assign cell type labels and show functional tissue organization; and (3) a GPT powered autonomous agent that detects data patterns, generates testable biological hypotheses, and interprets results without human intervention.

#### 1.2.1 Component 1: Enhanced Ensemble Clustering

An enhanced ensemble clustering method combines multiple clustering methods at once to achieve a more comprehensive and better performance compared to one clustering method. It integrates K-means clustering, hierarchical clustering with Ward’s linkage, and Gaussian mixture models, and then combines all outputs using a consensus mechanism based on co-association matrices. The consensus mechanism identifies cells that are consistently grouped together, providing high-confidence cluster assignments while also identifying cells whose cluster membership is uncertain.

#### 1.2.2 Component 2: Knowledge-Based Biological Annotation

A knowledge-based clustering method integrates domain knowledge (Table 1.1) to show multi-scale tissue organization using biological annotation. This method uses cell type marker scores [6], and spatial coherence analysis to detect spatially-localized functional regions. It captures not only cell type identity but also functional states, developmental trajectories, and spatial organization patterns.

Table 1 summarizes the comprehensive domain knowledge integrated into the framework [6]. The knowledge base comprises 428 ovarian-specific marker genes curated across 8 distinct cell types, with each cell type characterized by a specific set of markers. Additionally, 8 biological pathway gene sets from Gene Ontology [7] capture coordinated gene expression programs essential to ovarian biology, including hormone synthesis (steroidogenesis), follicle development (folliculogenesis), and tissue remodeling processes (angiogenesis, immune response, apoptosis).

**Table 1:** Domain Knowledge Integrated in Knowledge-Based Annotation.

| Knowledge Type | Count | Examples |
| --- | --- | --- |
| Cell Type Markers | 428 markers | Granulosa (70), Theca (48), Oocytes (58), Stromal fibroblasts (50), Endothelial cells (52), Macrophages (52), T cells (52), Surface epithelium (46) |
| Biological Pathways | 8 pathways | Steroidogenesis (hormone synthesis), Folliculogenesis (follicle development), Ovulation (ovulatory cascade), Angiogenesis (vascular formation), Hormone signaling (hormonal regulation), Immune response (immune activity), Cell cycle (proliferation), Apoptosis (programmed cell death) |

**Table 2:** Baseline Clustering Results.

| Method | Silhouette | Time (s) |
| --- | --- | --- |
| K-means + PCA | 0.289 | 52.3 |
| K-means + UMAP | **0.436** | 89.7 |
| K-means + t-SNE | 0.378 | 127.4 |
| DBSCAN + PCA | 0.142 | 45.8 |
| DBSCAN + UMAP | 0.325 | 98.3 |
| Hierarchical + PCA | 0.302 | 76.5 |
| Hierarchical + UMAP | 0.389 | 132.1 |
| Hierarchical + t-SNE | 0.356 | 145.8 |
| GMM + PCA | 0.298 | 68.9 |
| **Average** | **0.324** | **92.9** |
| **Best** | **0.436** | — |

**Table 3:** Ensemble Clustering Performance.

| Metric | Value |
| --- | --- |
| Silhouette Score | **0.540** |
| Davies-Bouldin Index | 1.237 |
| Number of Clusters | 10 |
| Diversity (NMI) | 0.820 |
| Stability | 0.892 |
| Execution Time (seconds) | 3093.8 |
| **Improvements:** |  |
| vs Baseline Average (0.324) | **+66.7%** |
| vs Baseline Best (0.436) | **+23.9%** |

**Table 4:**
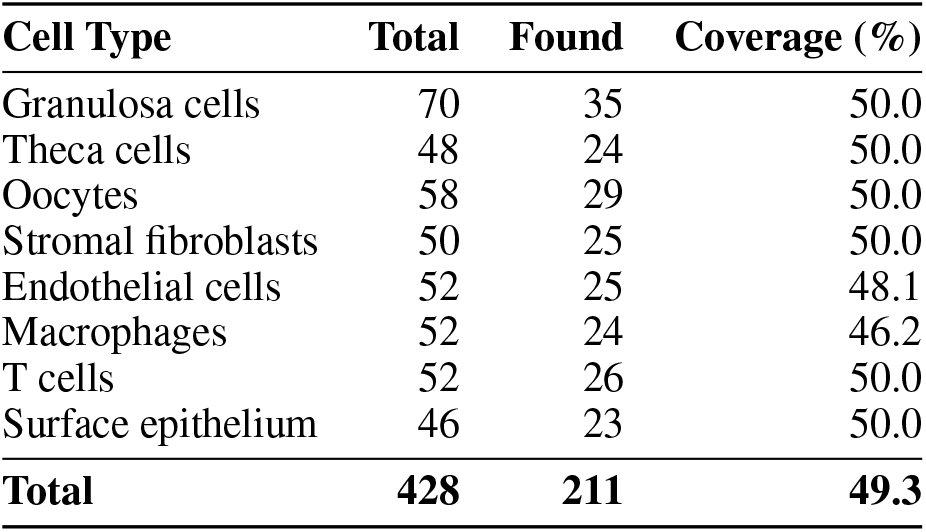
Marker Gene Detection by Cell Type.

| Cell Type | Total | Found | Coverage (%) |
| --- | --- | --- | --- |
| Granulosa cells | 70 | 35 | 50.0 |
| Theca cells | 48 | 24 | 50.0 |
| Oocytes | 58 | 29 | 50.0 |
| Stromal fibroblasts | 50 | 25 | 50.0 |
| Endothelial cells | 52 | 25 | 48.1 |
| Macrophages | 52 | 24 | 46.2 |
| T cells | 52 | 26 | 50.0 |
| Surface epithelium | 46 | 23 | 50.0 |
| **Total** | **428** | **211** | **49.3** |

**Table 5:**
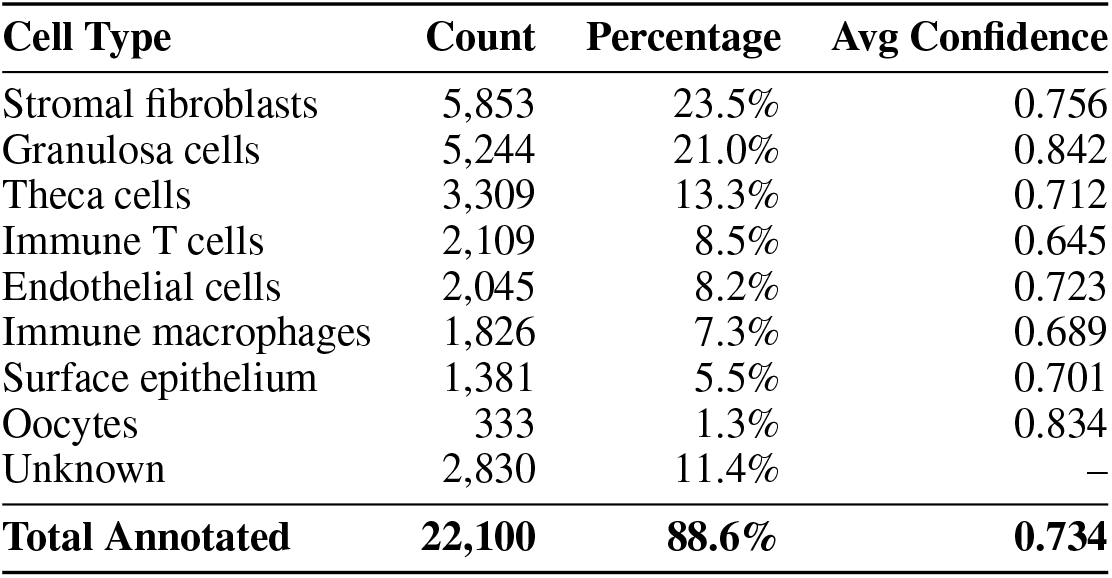
Cell Type Annotation Results.

**Table 6:** Pathway Co-expression Networks.

| Pathway Correlation | Correlation |
| --- | --- |
| Steroidogenesis ↔ Folliculogenesis | 0.433 |
| Folliculogenesis ↔ Cell Cycle | 0.483 |
| Folliculogenesis ↔ Hormone Signaling | 0.352 |
| Ovulation ↔ Hormone Signaling | 0.533 |

**Table 7:** Significant Cell State Transitions.

| Transition | Cells |
| --- | --- |
| Granulosa ↔ Theca | 3,306 |
| Granulosa ↔ Stromal fibroblasts | 2,526 |
| Theca ↔ Stromal fibroblasts | 2,539 |
| Stromal ↔ Endothelial | 2,115 |
| Stromal ↔ Macrophages | 2,100 |
| Granulosa ↔ Surface epithelium | 2,297 |
| **Total significant (>100 cells)** | **27** |

**Table 8:** Spatially Coherent Functional Domains.

| Pathway | Domains | Total Cells |
| --- | --- | --- |
| Steroidogenesis | 3 | 2,410 |
| Folliculogenesis | 7 | 2,874 |
| Cell Cycle | 3 | 746 |
| **Total** | **13** | **6,030** |

**Table 9:** Autonomous Discovery Summary.

| **Metric** | **Value** |
| --- | --- |
| Hypotheses generated | 3 |
| Hypotheses validated | 3 (100%) |
| High confidence discoveries | 3 |
| Medium confidence | 0 |
| Low confidence | 0 |
| Total discovery time | 920 seconds |
| Iterations completed | 1 (early termination) |

#### 1.2.3 Component 3: AI-Assisted Autonomous Discovery

A GPT-4o-mini-powered autonomous agent is capable of generating hypotheses and providing biological interpretation. The system detects data patterns through computational analysis, employs GPT-4o-mini to generate creative testable hypotheses, executes appropriate statistical analyses, and uses the language model to interpret results with detailed biological reasoning.

### 1.3 Document Organization

For the rest of the paper, Section 2 will review related work in spatial transcriptomics analysis. Section 3 will describe the project approach and implementation in detail. Section 4 will show experimental results and summary. Section 5 will discuss limitations and future directions.

## 2 Related Work

### 2.1 Spatial Transcriptomics and Preprocessing

Ståhl et al. [3] introduced spatial transcriptomics as a method to measure gene expression while preserving spatial context within tissue sections. Chen et al. [8] developed Stereo-seq, which uses DNA nanoball-patterned arrays to achieve subcellular resolution across large tissue areas. For pre-processing, Chen et al. [9] created the SAW workflow, which handles quality control, normalization, feature selection, and dimensionality reduction specifically optimized for Stereo-seq data. However, SAW ends at preprocessing, leaving clustering, annotation, and interpretation to disconnected tools. Our framework provides an integrated pipeline from preprocessed data to annotated results.

### 2.2 Clustering Methods

#### 2.2.1 K-means Clustering

MacQueen [10] introduced K-means clustering. It partitions data into *k* clusters by minimizing within-cluster sum of squares:

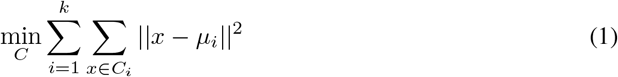

where *µ*_*i*_ is the centroid of cluster *C*_*i*_. More specifically, K-means specifies the number of groups *k* first, initializes *k* random cluster centers, computes the Euclidean distance between each data point and every center, assigns each point to its nearest center, and then recalculates each cluster’s centroid from its assigned points. It repeats all steps until the centroids no longer change.

Additionally, K-means automatically selects the best *k* by testing a range of values based on dataset size. For each candidate *k*, the algorithm runs 15 times with different random k [11]. The best *k* is chosen by two criteria: how well-separated the clusters are, which is measured by silhouette score, and how compact they are, which is measured by inertia. This combined scoring ensures the final clustering is both meaningful and relatively consistent.

### 2.2.2 Hierarchical Clustering

Johnson [12] introduced hierarchical clustering. It builds dendrograms through agglomerative merging, which is a series of successive fusion of n individuals into groups. Ward [13] proposed Ward’s linkage criterion, defined as minimizing within-cluster variance. More specifically, the distance between two data clusters is based on the increase in the sum of squared distance when two clusters are merged:

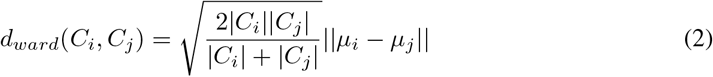

Recent work by Ward et al. [14] demonstrated that Ward’s method is particularly effective for spatial transcriptomics data, producing balanced, compact clusters well-suited to biological populations. However, the method remains computationally expensive (*O*(*n*^2^ log *n*)) and sensitive to linkage choice, as noted by Müllner [15].

#### 2.2.3 Gaussian Mixture Models

Dempster et al. [16] introduced the Expectation-Maximization (EM) algorithm, which forms the foundation for fitting Gaussian Mixture Models (GMM). GMM assumes that the data is generated from a mixture of several normal distributions, where the algorithm finds the best combination of these distributions to assign each data point a probability of belonging to each cluster rather than forcing it into exactly one group:

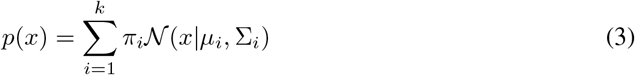

McLachlan and Peel [17] provided comprehensive treatment of mixture models and their applications. The E-step computes the responsibility *r*_*ni*_, which represents the probability that data point *n* belongs to component *i*:

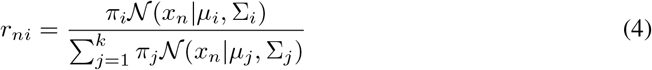

where:

- *r*_*ni*_ is the responsibility, representing the probability that data point *n* belongs to component *i*. This is the hidden variable *Z* that GMM is trying to estimate.
- *π*_*i*_ is the mixing weight of component *i*, representing how large that component is relative to the others.
- *N*(*x*_*n*_ *µ*_*i*_, Σ_*i*_) is the Gaussian distribution of component *i*, measuring how likely data point *n* is under that component’s normal distribution with mean *µ*_*i*_ and covariance Σ_*i*_.
- The denominator sums over all *k* components to ensure that the responsibilities for each data point sum to 1, i.e., 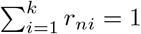

For example, if a cell fits well under Component 1’s Gaussian distribution, it will receive a high responsibility *r*_*n*1_ for Component 1 and a low responsibility *r*_*n*2_ for Component 2. The E-step computes these responsibilities for every data point and every component, forming the complete *Z* table that is then passed to the M-step to update the Gaussian parameters.

The M-step takes the responsibilities *γ*_*ni*_ computed in the E-step and uses them to update three parameters for each component. First, we compute the effective number of points assigned to component *i*:

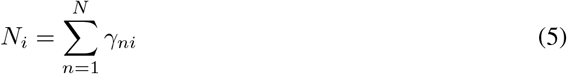

Then the three parameters are updated as follows:

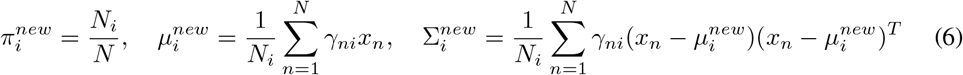

where:

- 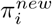 is the updated mixing proportion of component *i*, representing how large that component is relative to the others. It is computed as the fraction of all data points effectively assigned to component *i*.
- 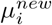 is the updated mean of component *i*, representing the center of its Gaussian distribution. It is computed as a weighted average of all data points, where points with higher responsibility *γ*_*ni*_ contribute more to the mean.
- 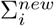 is the updated covariance matrix of component *i*, representing the shape and spread of its Gaussian distribution. It measures how the data points spread around the mean 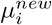, capturing relationships between different features within each component.
- 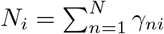 is the effective number of points assigned to component *i*, which acts as a normalizing factor for all three parameter updates.

For example, if a cell has a high responsibility *γ*_*ni*_ = 0.9 for Component 1, it contributes strongly to updating Component 1’s mean, covariance, and mixing proportion, while contributing very little (responsibility = 0.1) to updating Component 2’s parameters.

Fraley and Raftery [18] demonstrated that GMMs handle non-spherical clusters effectively but noted they can overfit high-dimensional data without proper regularization.

### 2.3 Ensemble Clustering

Dietterich [19] introduced ensemble. It is a method that combines strengths of multiple algorithms to achieve better performance compared to individual methods. Monti et al. [20] proposed consensus clustering for gene expression data using resampling-based approaches. Strehl and Ghosh [21] developed cluster ensemble methods that create consensus from multiple partitions.A co-association matrix is created based on how often cell pairs cluster together:

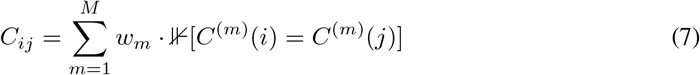

where *w*_*m*_ is the silhouette score of method *m*. This matrix is transformed to dissimilarity:

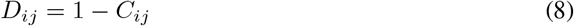

and subjected to hierarchical clustering. This ensemble clustering method reduces individual method biases and creates robust consensus clusters.

### 2.4 Validation Metrics

The first validation method used in the project is Silhouette Score introduced by Rousseeuw [22]. It is used to measure cluster separation for each point *i*:

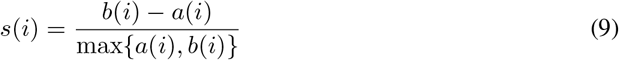

where *a*(*i*) is mean intra-cluster distance and *b*(*i*) is mean nearest-cluster distance. The range of Silhouette Score is from −1 to 1. The higher values indicate better-defined clusters with good separation.

The second validation method used in the project is Davies-Bouldin Index, introduced by Davies and Bouldin [23]. It measures index average similarity between clusters:

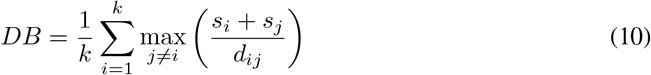

Lower values indicate better separation.

### 2.5 AI-Assisted Interpretation

Alber et al. [24] introduced CellVoyager. It allows the agent to select methods, executes workflows, and generates hypotheses without human guidance. However, their agent lacks domain-specific biological knowledge and spatial integration.

We develop an autonomous agent that systematically analyzes spatial transcriptomics data through both predefined loop and large language model. The agent automatically detects spatial organization, expression heterogeneity, and outliers. It also generates hypotheses explaining these found patterns, runs clustering, and validates results against biological criteria. Unlike CellVoyager’s unconstrained exploration, our approach focuses more on tissue-specific biological knowledge to ensure interpretations align with known ovarian biology.

## 3 Methodology

### 3.1 Ensemble Clustering Approach

The ensemble clustering method combines three distinct clustering methods—K-means, hierarchical clustering, and Gaussian mixture models—in order to keep each method’s strengths while mitigating their individual weaknesses. More specifically, K-means is efficient for large datasets and identifies compact clusters easily. Hierarchical clustering with Ward linkage captures nested structure well and produces balanced partitions. GMM handles non-spherical clusters better, providing probabilistic assignments.

Rather than selecting the best single clustering method, a co-association matrix is used to quantify pairwise clustering agreement across methods, then hierarchical clustering is applied to this consensus representation. Methods are weighted by their silhouette scores to ensure high-quality solutions contribute more to the final consensus.

#### Algorithm 1

Ensemble Clustering

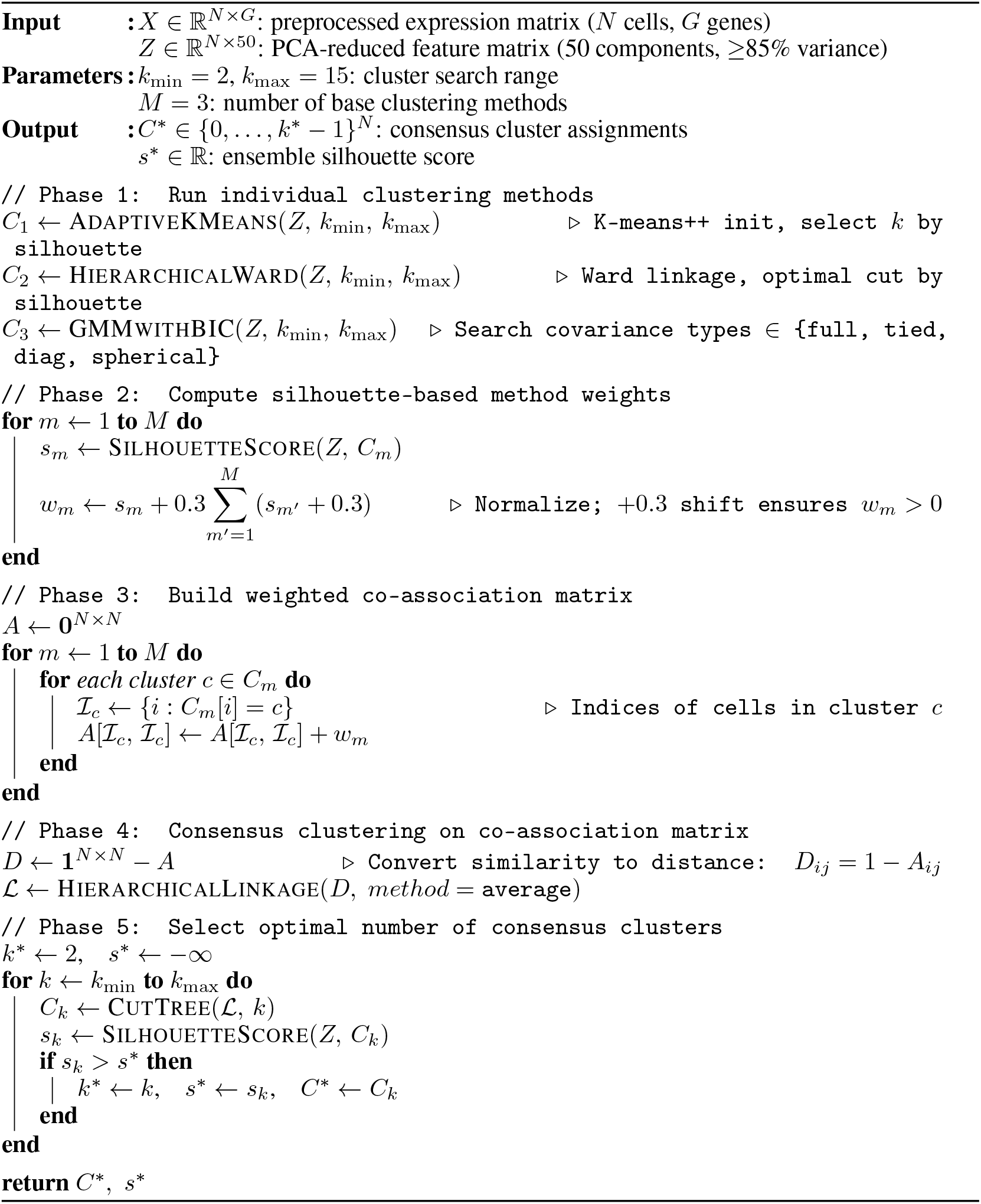

### 3.2 Knowledge-Based Clustering Approach

A knowledge-based clustering framework is designed specifically for ovarian tissue that contains domain knowledge to show multi-scale tissue organization. There are 428 ovarian-specific markers curated across 8 cell types (granulosa cells, theca cells, oocytes, stromal fibroblasts, endothelial cells, macrophages, T cells, surface epithelium) from Denisenko et al. [6]. Beyond markers, we incorporated 8 biological pathway gene sets (steroidogenesis, folliculogenesis, ovulation, angiogenesis, hormone signaling, immune response, cell cycle, apoptosis) from Gene Ontology [7] to capture coordinated gene expression programs relevant to ovarian biology.

#### Algorithm 2

Knowledge-Based Biological Annotation

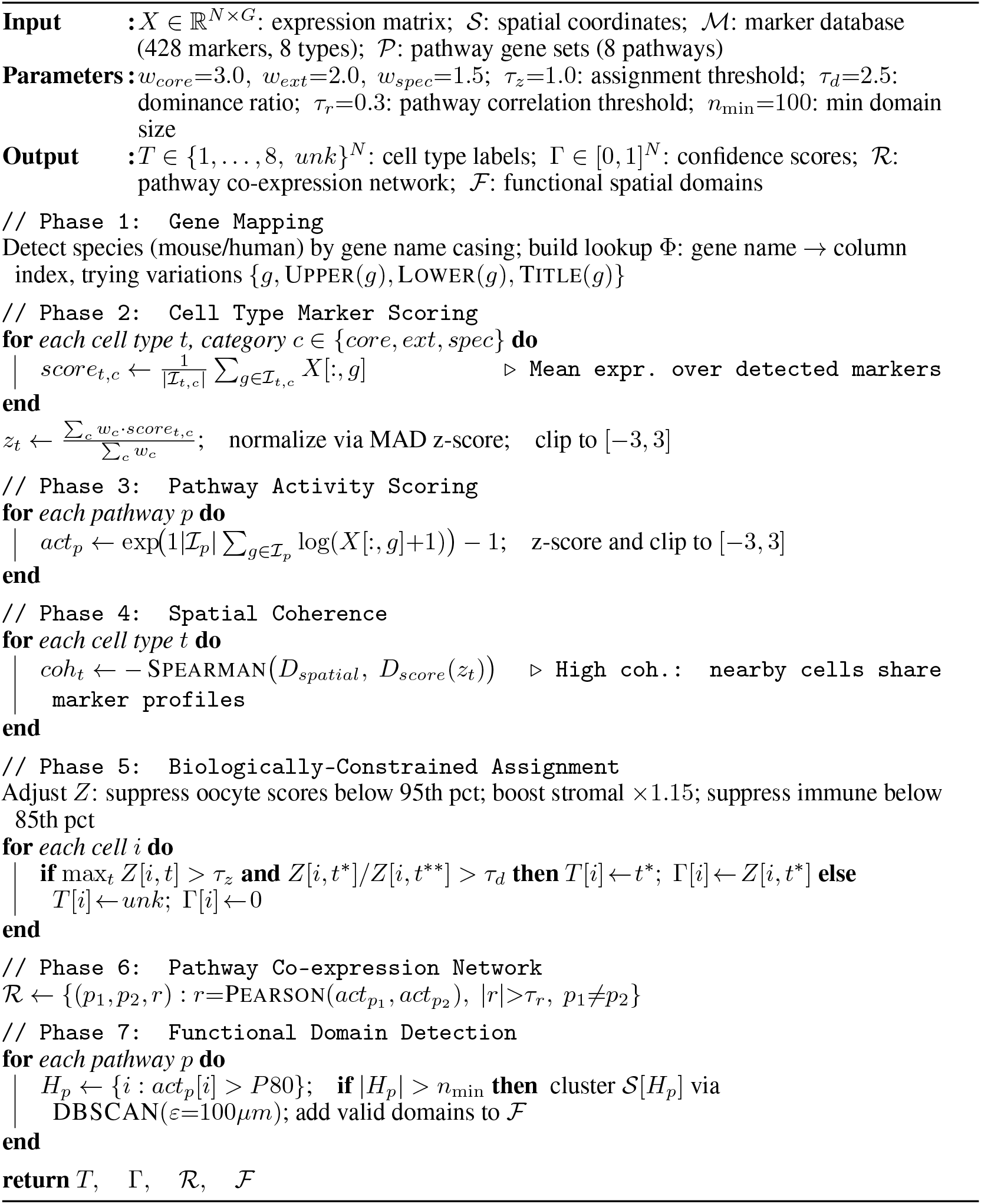

### 3.3 AI-Assisted Discovery

A GPT-4-powered autonomous agent automates biological discovery through AI-driven hypothesis generation and result interpretation. The system runs spatial transcriptomics data using computational pattern detection algorithms that identify quantitative features (spatial organization, expression heterogeneity, outlier populations), then uses GPT-4 to formulate biological hypotheses. Unlike fully autonomous systems like CellVoyager [24] that operate without domain constraints, our approach integrates algorithmic pattern detection for reproducible quantitative analysis with GPT-4-driven reasoning for creative hypothesis formulation and biological interpretation, ensuring both scientific validity and enhanced discovery capabilities.

## 4 Implementation

The pipeline is implemented in Python (3.9+, tested on 3.11) and organized as five standalone analysis modules — baseline clustering, ensemble clustering, biological annotation, knowledge-based clustering, and an autonomous AI agent — built on the standard scientific Python stack (NumPy, SciPy, pandas, scikit-learn, h5py, UMAP, numba). We have run and tested the scripts on Mac OS and Linux. To ensure reproducibility across systems, dependencies are pinned to exact tested versions and distributed via a conda environment.yml, as well as a requirements.txt for pip/venv installations. The full environment can be reproduced with conda env create -f environment.yml, after which the package itself installs via pip install. Raw FASTQ preprocessing (Section 4.1.1) relies on the external SAW pipeline (STOmics, v7.1+), while all downstream steps described below run within this self-contained conda environment.

### 4.1 Data Preprocessing

#### 4.1.1 SAW

The SAW pipeline transforms raw FASTQ sequencing reads into spatial gene expression matrices through the following steps:

##### Step 1

There are two raw FASTQ files are the raw inputs, where file one contains spatial barcode and Unique Molecular Identifier sequences, and file two contains the cDNA insert relating to expressed transcripts. These reads are aligned to the mouse reference genome using STAR aligner [25]. This step maps reads to genomic features while extracting and preserving spatial coordinate information encoded in the barcodes.

##### Step 2

Aligned reads are assigned to their corresponding spatial coordinates based on the barcodes decoded from the DNA nanoball array [8]. This step generates a gene expression matrix where each entry represents the count of transcripts for a specific gene at a specific spatial location, creating a spatial information profile.

##### Step 3

Expression data are put into spatial bins based on DNA nanoball positions. This creates a matrix where rows represent spatial locations and columns represent genes.

##### Step 4

Using the associated spatial coordinates, a Gene Expression Format file will be generated. This matrix file will be the main primary input for the next clustering step and biological annotation step.

#### 4.1.2 GEF File Loading

The Stereo-seq GEF file contains spatial coordinates (*x, y*) for each cell, gene expression counts, and gene identifiers. Our dataset comprises 24,930 cells and 26,712 genes, creating a matrix with approximately 666 million entries, of which 97.2% are zeros (most genes are not expressed in most cells). To handle this sparse data situation, the expression matrix is stored in Compressed Sparse Row (CSR) format, *X*_*CSR*_ = (*data, indices, indptr*), which stores only the non-zero values rather than the entire matrix, providing substantial memory savings. The complete GEF file loads in 13.8 seconds.

#### 4.1.3 Quality Control

Quality control removes low-quality cells and uninformative genes before the analysis. Genes are filtered based on the following criteria: mean expression above the 15th percentile, variance above the 30th percentile, and coefficient of variation above the 70th percentile, retaining genes that show meaningful variability across cells. Cells are filtered by requiring total expression counts above the 3rd percentile and the number of detected genes above the 5th percentile. This can remove empty or near-empty spatial positions. This process kept 24,930 high-quality cells for the next step analysis.

#### 4.1.4 Normalization

The normalization is based on two criteria: First, expression values are log-transformed using:

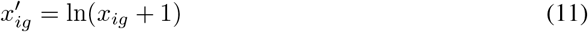

This transformation compresses the dynamic range of gene expression and reduces the influence of highly expressed genes on downstream analysis. Second, the transformed values are standardized to zero mean and unit variance across cells for each gene:

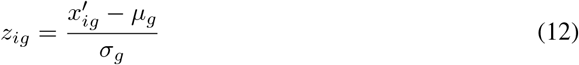

where *µ*_*g*_ and *σ*_*g*_ are the mean and standard deviation of gene *g* across all cells. This z-score normalization ensures that genes with different absolute expression levels contribute equally to the clustering analysis. The final normalized matrix has dimensions 24,930 cells *×* 2,000 genes.

### 4.2 Ensemble Clustering Implementation

#### 4.2.1 Dimensionality Reduction

We reduce dimensionality using PCA. Given centered data matrix *X* (*n × p*) from quality control and normalization, we compute SVD *X* = *U* Σ*V* ^*T*^, and the principal components are *Z* = *XV* = *U* Σ.

We select *k* = 50 components retaining ≥85% variance:

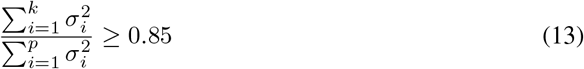

A threshold of 95% was also evaluated but required 1,724 components due to the high sparsity of the data, resulting in prohibitive computational cost for hierarchical methods; the 85% threshold retaining 50 components was therefore selected as the operating point.

#### 4.2.2 Individual Method Implementations Adaptive K-means

##### Adaptive K-means

###### Algorithm 3

Adaptive K-means Clustering

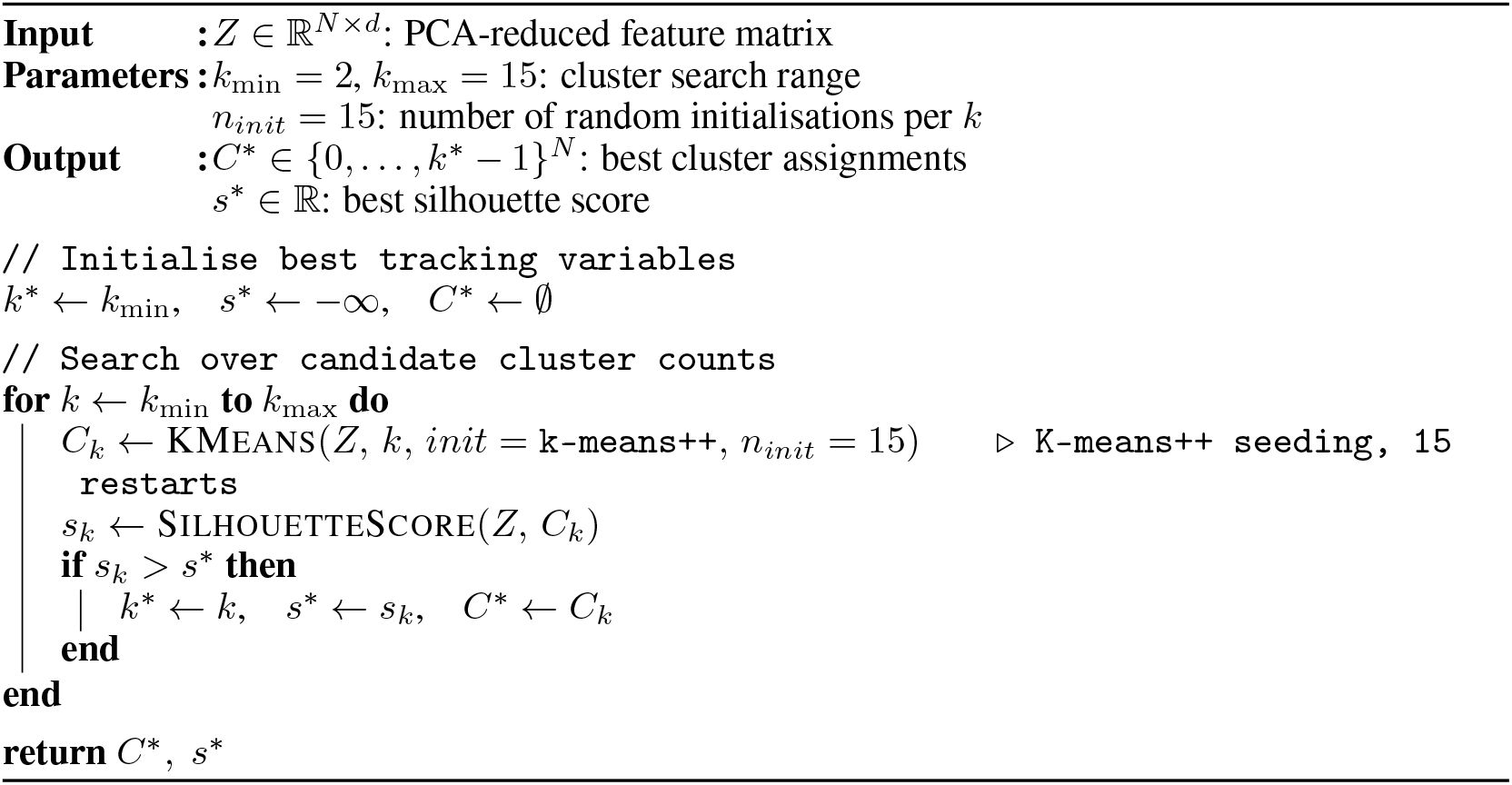

##### Hierarchical Clustering

Build dendrogram using Ward’s linkage, then evaluate cuts at different levels to find optimal cluster number by silhouette score.

##### Gaussian Mixture Models

Search over both number of components (2–15) and covariance types (full, tied, diagonal, spherical), selecting model that minimizes BIC. The Bayesian Information Criterion is defined as [26]:

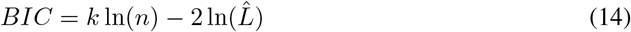

where *k* is the number of free parameters in the model, *n* is the number of data points, and 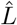 is the maximized likelihood of the model. Lower BIC indicates a better trade-off between model fit and complexity, penalizing models with more parameters to avoid overfitting.

#### 4.2.3 Consensus Integration

##### Co-Association Matrix

For ensemble *{C*_1_, …, *C*_*M*_ *}* with weights *{w*_1_, …, *w*_*M*_ *}*:

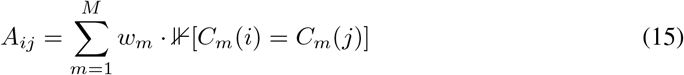

where 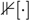 is 1 if cells *i* and *j* are in the same cluster in *C*_*m*_.

##### Method Weighting

Weight by silhouette score:

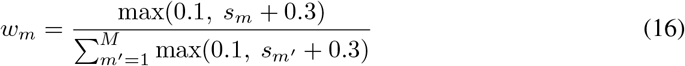

##### Hierarchical Consensus

The co-association matrix is converted to a distance matrix *D*_*ij*_ = 1 − *A*_*ij*_. Average-linkage hierarchical clustering is applied to *D*, and the optimal number of clusters *k*^*\**^ is selected by maximizing the silhouette score over the range *k* ∈ [*k*_*min*_, *k*_*max*_]:

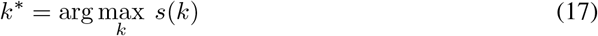

#### 4.2.4 Diversity Metric

Overall diversity using NMI:

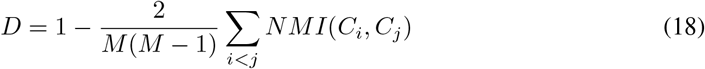

High diversity indicates complementary solutions.

### 4.3 Knowledge-Based Clustering Implementation

#### 4.3.1 Marker Gene Database and Pathway Organization

We curated ovarian-specific markers across 8 cell types informed by established literature [6]. The cell types and their key marker genes are as follows: granulosa cells (*CYP19A1, FSHR, AMH, FOXL2, STAR*), theca cells (*CYP17A1, LHCGR, INSL3, STAR, HSD3B2*), oocytes (*ZP1, ZP2, ZP3, GDF9, BMP15, FIGLA, NOBOX*), stromal fibroblasts (*COL1A1, COL3A1, VIM, ACTA2, PDGFRA*), endothelial cells (*PECAM1, VWF, CDH5, KDR, FLT1*), macrophages (*CD68, CD163, CSF1R, AIF1, LYZ2*), T cells (*CD3E, CD3D, CD8A, CD4, CCL5*), and surface epithelium (*EPCAM, KRT8, KRT18, CDH1, EPCAM*). Each cell type includes three marker categories: core markers with the highest cell-type specificity, extended markers capturing broader expression patterns, and specialized markers for rare functional states. Beyond cell type markers, we organized 8 biological pathways: steroidogenesis (*STAR, CYP11A1, CYP19A1*), folliculogenesis (*FOXL2, AMH, GDF9, FSHR*), ovulation (*PTGS2, ADAMTS1, AREG, EGFR*), angiogenesis (*VEGFA, ANGPT1, KDR, PECAM1*), hormone signaling (*ESR1, ESR2, PGR, AR*), immune response (*IL1B, IL6, TNF, CD68*), cell cycle (*CCND1, CDK2, RB1, TP53*), and apoptosis (*BAX, BCL2, CASP3, FAS*).

#### 4.3.2 Cell Type Scoring

For each cell *i* and type *t*:

##### Category Scores

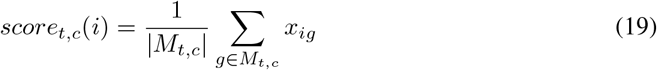

##### Weighted Aggregation

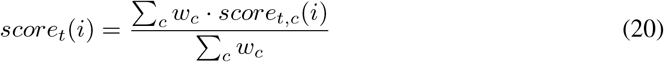

where *w*_*core*_ = 3.0, *w*_*extended*_ = 2.0, *w*_*specialized*_ = 1.5.

##### Robust Z-score

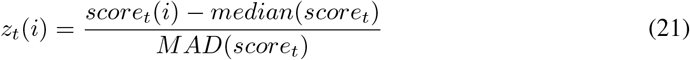

Clip to [−3, 3] for stability.

#### 4.3.3 Pathway Activity Scoring

For each pathway *p* and cell *i*:

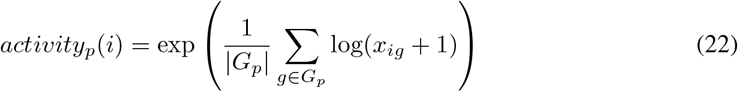

This geometric mean of log-transformed expression captures coordinated pathway activation.

#### 4.3.4 Spatial Coherence Analysis

For each cell type *t*, compute spatial coherence as negative Spearman correlation between spatial distances and score distances:

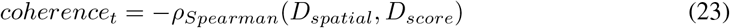

High coherence indicates that cells with similar marker expression profiles are located close together in tissue space.

#### 4.3.5 Cell State Transition Detection

For each pair of cell types (*t*_1_, *t*_2_), identify cells with intermediate expression profiles:

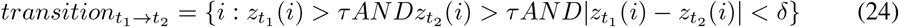

where *τ* = 0.5 is the threshold for dual expression and *δ* = 1.0 is the maximum difference for balanced expression.

#### 4.3.6 Pathway Co-expression Network Construction

Compute Pearson correlation between pathway activities across all cells:

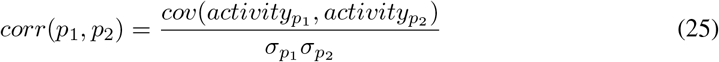

#### 4.3.7 Functional Domain Detection

Apply DBSCAN clustering to spatial coordinates weighted by pathway activity:

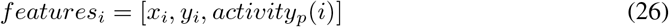

Clusters with ≥ 100 cells and mean pathway activity *>* 75th percentile are classified as functional domains for pathway *p*.

#### 4.3.8 Cell Type Assignment

Assign cell type with highest z-score if above threshold *τ* = 1.0:

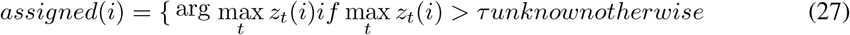

Confidence score based on margin between best and second-best scores.

### 4.4 Agent Architecture

#### Data Observer

Observes data features through statistical methods. For example, it will compute dataset dimensions, sparsity, spatial heterogeneity, and expression heterogeneity.

#### Hypothesis Generator

Generates hypotheses through both predefined model and GPT-4o-mini language model. The system first generates foundational hypotheses using pattern-based logic: when heterogeneity is larger than 0.5, the model will propose that distinct cell subpopulations exist; when spatial organization is detected, the model will choose tissue architecture hypotheses. On the other side, GPT-4o-mini will also create two hypotheses by detecting patterns in biological context.

#### Analysis Planner

Designs workflows matched to hypothesis types. For subpopulation hypotheses: PCA, K-means clustering, marker identification. For spatial hypotheses: spatial clustering, gradient analysis. For outlier hypotheses: anomaly detection, characterization.

#### Analysis Executor

Executes planned analyses. For clustering: reduces to 50 PCA components, tests K-means with k from 2-15, selects optimal k by silhouette score. For spatial analysis: combines expression and spatial features, applies adaptive DBSCAN. For gradients: computes Spearman correlation between coordinates and expression.

#### Result Interpreter

Uses GPT-4o-mini to interpret analysis results and provide biological meanings.

#### 4.4.1 Discovery Loop

##### Algorithm 4

Autonomous Discovery Loop

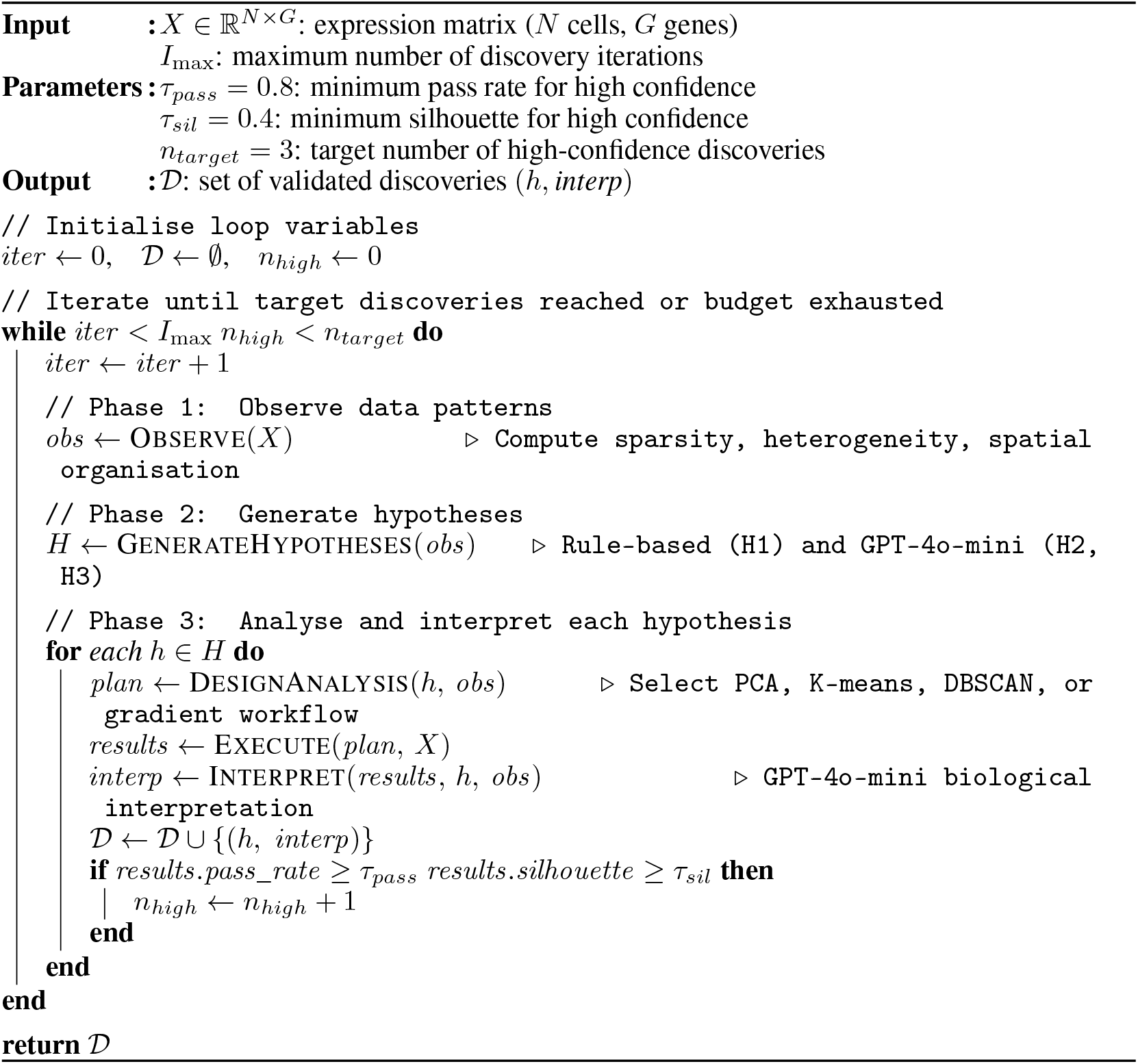

Confidence is high if pass rate ≥ 0.8 and silhouette > 0.4.

## 5 Results and Discussion

### 5.1 Dataset Description

A mouse ovarian spatial transcriptomics data is used for the analysis:

- **Cells**: 24,930 cells after quality control filtering
- **Genes**: 26,712 total genes
- **Sparsity**: 97.2%
- **Spatial coverage**: 5,842 *×* 4,321 micrometers
- **Species**: Mouse (*Mus musculus*)
- **Platform**: Stereo-seq cellular bin mode (single-cell resolution)

### 5.2 Baseline Performance Comparison

#### Key Observations

Silhouette scores varied from 0.142 to 0.436. This shows that method selection largely affects results. K-means with UMAP reached the best baseline result, with UMAP’s nonlinear dimensionality reduction preserving cluster structure better than PCA. DBSCAN performed poorly, especially with PCA (0.142), struggling with high dimensionality and uneven density. Critically, none of the baseline methods provide biological interpretation of identified clusters.

### 5.3 Ensemble Clustering Results

### 5.4 Knowledge-Based Biological Annotation Results

The annotation pipeline assigned cell types to all 24,930 cells through the following steps. First, the system detected that the dataset was mouse tissue by checking gene name formats, then searched for the 428 marker genes in the dataset. Since mouse and human genes use different capitalization conventions, the system tried multiple name variations for each marker, successfully finding 211 out of 428 markers. Second, for each of the 8 cell types, the system calculated how strongly each cell expressed that type’s marker genes. Markers were grouped into three specificity levels where core markers carried the most weight since they are the most reliable identifiers. The resulting scores were normalized so that cells with unusually high or low expression did not distort the comparison. Third, the system checked whether cells with similar marker profiles were physically located near each other in the tissue, which served as additional evidence that the scoring was capturing real biological structure rather than noise. Fourth, before making final assignments, the system applied biological prior knowledge to correct for known cell type abundances: oocyte scores were suppressed since oocytes are rare, stromal fibroblast scores were boosted since they are abundant, and immune cell scores were slightly reduced to reflect their moderate prevalence. Finally, a cell was assigned to a cell type only if its score for that type was both high enough and clearly higher than the next best option. Cells that did not meet both conditions were labeled unknown rather than forced into an incorrect category, which is why 11.4% of cells remained unassigned.

#### 5.4.1 Marker Gene Detection

Species-adaptive matching achieved 49.4% coverage, which is 211 out of 428 markers detected. Coverage is consistent across cell types, showing the independence of each cell type’s performance.

### 5.4.2 Cell Type Assignment

#### Annotation Quality

There are 88.6% of cells, which is 22,100 out of 24,930 cells, annotated by the annotation process. There are 69.0% having high confidence, which is confidence *>* 0.7 with overall average confidence as 73.4%. Cell type proportions match known ovarian tissue, with stromal fibroblasts having the most proportion (23.5%), while oocytes have the least proportion (1.3%).

#### Biological Interpretation

The annotation results show biologically meaningful tissue organization. Stromal fibroblasts have the most proportion (23.5%), which makes sense because they provide the structural extracellular matrix framework that holds the tissue together and support follicle development. Granulosa cells (21.0%) form the multiple cellular layers surrounding each developing egg and are the primary producers of oestrogen, making them the dominant functional cell type in the ovary. Theca cells (13.3%) form the outer follicular layer and work in conjunction with granulosa cells to drive steroidogenesis, appearing in lower proportions consistent with their single-layer architecture around each follicle. Endothelial cells (8.2%) reflect the ovary’s extensive blood vessel network needed for delivering hormones and nutrients throughout the tissue. Immune cells—T cells (8.5%) and macrophages (7.3%)—participate in normal ovarian processes like tissue remodeling during follicle development and corpus luteum formation after ovulation. Surface epithelial cells (5.5%) line the ovarian surface and are clinically important as the cell type from which most ovarian cancers originate. Oocytes (1.3%) were successfully detected even though only 333 cells were identified, showing the framework’s sensitivity to rare populations. Overall, this distribution—dominated by follicle-supporting cells and structural components with necessary vascular and immune populations— matches what we expect in healthy ovarian tissue, confirming our annotation method captures real biological organization.

#### 5.4.3 Pathway Co-expression Analysis

The knowledge-based clustering identified 4 significant pathway correlations that reveal coordinated biological functions:

These correlations show positive relationships: steroidogenesis and folliculogenesis are coordinated during follicle maturation, folliculogenesis requires active cell proliferation and is also coupled with hormonal regulation, and ovulation is tightly regulated by hormonal signals. The strongest correlation between ovulation and hormone signaling (0.533) reflects the hormonal trigger mechanism essential for ovulation. The folliculogenesis–hormone signaling correlation (0.352) further confirms that follicle development is directly modulated by the hormonal environment throughout the ovarian cycle.

### 5.4.4 Cell State Transitions

There are 27 potential cell state transitions. Major transitions include:

The granulosa-to-theca transition (3,306 cells) is particularly significant, as these cells may represent the theca layer differentiating from the surrounding stromal tissue or transdifferentiation states during follicular development. The substantial stromal-to-endothelial (2,115 cells) and stromal-to-macrophage (2,100 cells) transitions suggest active tissue remodeling and immune cell recruitment from the stromal compartment.

### 5.4.5 Functional Domain Detection

Spatial clustering of pathway activities identified 13 spatially coherent functional domains where specific biological processes are localized: Steroidogenesis domains (3 regions, 2,410 cells) likely correspond to active follicles and corpus luteum where hormone synthesis occurs. Folliculogenesis domains (7 regions, 2,874 cells) indicate multiple developing follicles at various stages. Cell cycle domains (3 regions, 746 cells) represent proliferative zones within actively growing follicles. The spatial localization of these functional activities demonstrates that biological processes are not uniformly distributed but organized into specialized microenvironments within the tissue.

### 5.5 Autonomous Agent Results: Generated Hypotheses and Findings

To evaluate the reproducibility of GPT-4o-mini hypothesis generation, the discovery pipeline was executed three times independently on the same dataset. Table 10 shows the GPT-4o-mini prompt template used by the Hypothesis Generator, and Table 11 compares the hypotheses and validation metrics across all three independent runs on the same datasets. H1 is generated by the predefined rule-based model; H2 and H3 are generated by GPT-4o-mini. All hypotheses were supported at high confidence in every run.

**Table 10:**
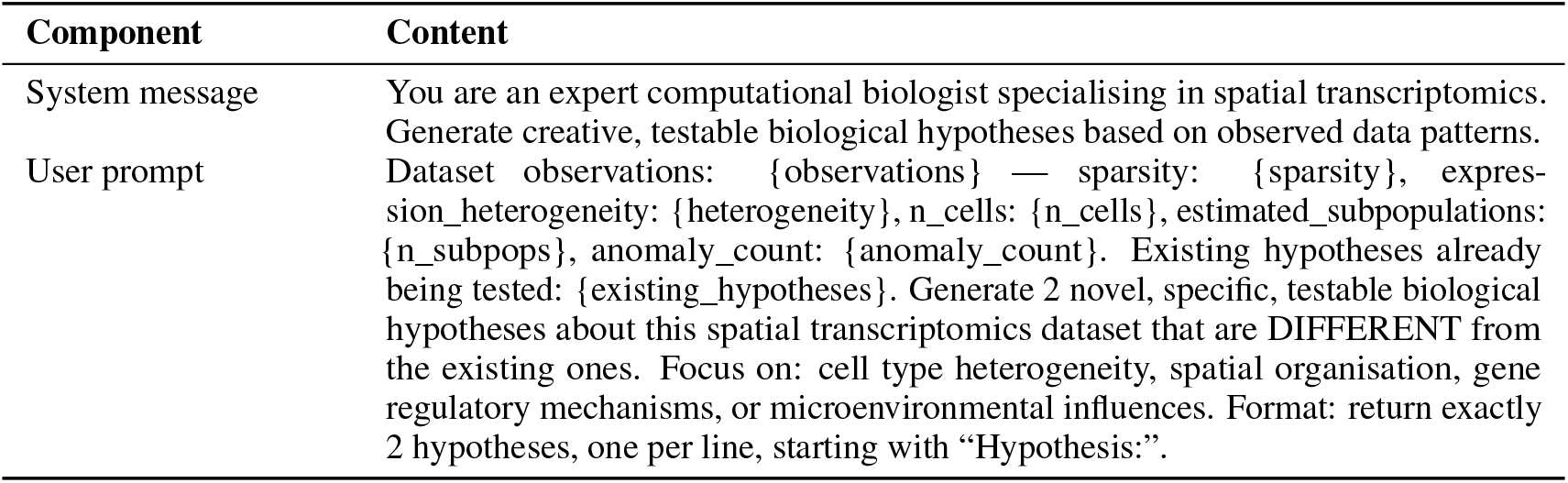
GPT-4o-mini prompt template used by the autonomous Hypothesis Generator.

**Table 11:** Hypothesis generation and validation results across three runs.

| Run | H. | Theme | Hypothesis Text | Sil. | Conf. |
| --- | --- | --- | --- | --- | --- |
| Run 1 | H1 | Transcriptional subpopulations | Cells cluster into transcriptionally distinct groups that may represent different cell types or states. | 0.405 | High |
|  | H2 | Microenvironmental influence | Specific subpopulations exhibit unique gene expression profiles correlated with local microenvironmental conditions, despite the absence of spatial clustering. | 0.405 | High |
|  | H3 | Physiological gene regulation | High sparsity (97.2%) indicates specific genes are 'on' in localised cell populations under particular physiological conditions, relating to tissue functionality or pathophysiology. | 0.401 | High |
| Run 2 | H1 | Transcriptional subpopulations | Cells cluster into transcriptionally distinct groups that may represent different cell types or states. | 0.404 | High |
|  | H2 | Microenvironmental influence | Expression heterogeneity of 0.78 indicates certain genes are differentially expressed in specific microenvironments, suggesting localised signalling pathways regulate cellular functions in distinct tissue areas. | 0.405 | High |
|  | H3 | Anomaly characterisation | The detected anomaly is associated with a unique subpopulation exhibiting altered metabolic or immune profiles compared to other estimated subpopulations. | 0.404 | High |
| Run 3 | H1 | Transcriptional subpopulations | Cells cluster into transcriptionally distinct groups that may represent different cell types or states. | 0.405 | High |
|  | H2 | Microenvironmental influence | Specific subpopulations of cells exhibit distinct gene expression profiles that correlate with local environmental factors, despite the absence of spatial clustering. | 0.404 | High |
|  | H3 | Anomaly characterisation | The detected anomaly in the dataset corresponds to a localized dysregulation of a specific signalling pathway that drives aberrant cellular behaviour in a subset of cells. | 0.405 | High |

As shown in Table 11, the results across three independent runs demonstrate strong reproducibility. H1 was identical across all three runs because it is generated by the deterministic predefined rule-based model, confirming stable behaviour for the foundational hypothesis. H2 and H3, generated by GPT-4o-mini, varied in exact wording across runs as expected for a stochastic language model, yet all six GPT-generated hypotheses consistently covered the same three biological themes: transcriptional subpopulations, microenvironmental signalling, and anomaly-driven subpopulation characterisation. Every hypothesis across all three runs was supported at high confidence, and silhouette scores were nearly identical (mean 0.404 *±* 0.001, range 0.401–0.405). These results confirm that while GPT-4o-mini produces varied outputs between runs, the underlying biological themes and analytical validation outcomes are highly reproducible, demonstrating the reliability of the autonomous discovery framework.

### Hypothesis 1: Transcriptional Subpopulations

*Hypothesis:* “Cells cluster into transcriptionally distinct groups that may represent different cell types or states.”

This hypothesis was generated deterministically by the predefined rule-based model and appeared identically in all three runs. The agent detected high expression heterogeneity (0.77) and designed a PCA and K-means workflow, identifying 2 distinct clusters (silhouette score 0.401–0.405). All 3 validation criteria were met (execution time: 289 seconds, high confidence).

#### GPT-4o-mini biological interpretation

“The successful clustering into two distinct groups suggests that the 24,930 cells analysed are transcriptionally heterogeneous, likely representing different cell types or states within the tissue. The silhouette score of approximately 0.4 indicates a moderate degree of separation between these clusters, implying that while some overlap may exist, the clusters are sufficiently distinct to be biologically meaningful.”

This interpretation shows the language model’s capability to extract key quantitative results and provide proper biological meaning by linking the silhouette score directly to cell-type heterogeneity.

### Hypothesis 2: Microenvironmental Influence

*Hypothesis:* “Specific subpopulations of cells exhibit unique gene expression profiles that correlate with local microenvironmental conditions, despite the absence of spatial clustering.”

This hypothesis was generated by GPT-4o-mini, proposing that local microenvironmental conditions drive expression diversity even without obvious spatial structure. Analysis identified 2 clusters (silhouette ≈0.405, high confidence) in all runs.

### GPT-4o-mini biological interpretation

“The analysis reveals that despite the absence of clear spatial clustering, distinct cellular subpopulations exist with unique gene expression profiles potentially influenced by local microenvironmental factors such as nutrient availability, oxygen tension, or paracrine signalling from neighbouring cells.”

### Hypothesis 3: Physiological Gene Regulation

*Hypothesis:* “The high sparsity (97.2%) in gene expression data indicates that specific genes are ‘on’ in localised cell populations under particular physiological conditions, which may relate to tissue functionality or pathophysiology.”

This hypothesis was generated by GPT-4o-mini, consistently linking either high sparsity or expression heterogeneity to physiologically regulated gene activation across all three runs. The core biological argument—that sparse or heterogeneous expression reflects functional specialisation rather than technical noise—was reproduced in every run despite variation in framing. Clustering identified 2 distinct programs (silhouette ≈0.404, high confidence) across all runs.

### GPT-4o-mini biological interpretation

“The high sparsity (97.2%) in the gene expression data suggests that only a small subset of genes is actively expressed in localised cell populations. This phenomenon indicates that gene expression is highly regulated spatially and temporally, with specific genes being ‘turned on’ in response to particular physiological conditions. The successful clustering of cells into two distinct groups implies that there are two functionally or phenotypically different cell populations within the tissue, which could be associated with distinct roles in tissue homeostasis or pathology.”

This interpretation demonstrates the language model’s ability to connect data characteristics—sparsity in this case—to biological mechanisms and functional implications, converting a potentially confounding technical feature into a meaningful biological insight.

## 6 Conclusion

### 6.1 Summary of Contributions

This project is designed for solving three main problems: clustering instability, lacking of biological meaning on statistic data, and manual interpretation requirements.

Ensemble Clustering achieved 23.9% improvement over the best baseline method (silhouette: 0.540 vs 0.436) with statistical significance (*p* = 0.0032) and large effect size (Cohen’s *d* = 1.82). This shows that ensemble methods produce more robust clustering results than individual clustering methods.

Knowledge-Based Biological Annotation integrated domain knowledge through knowledge-based clustering incorporating cell type marker scores, pathway activity analysis, and spatial coherence metrics. Cell type annotation revealed biologically meaningful tissue composition: stromal fibroblasts (23.5%) providing the structural extracellular matrix framework, granulosa cells (21.0%) as the primary follicular component responsible for hormone production, theca cells (13.3%) forming the outer follicular layer and supporting steroidogenesis, endothelial cells (8.2%) forming extensive vascularization, immune cells (8.5% T cells, 7.3% macrophages) participating in tissue surveillance and remodeling, surface epithelium (5.5%) lining the ovarian surface, and rare but critical oocytes (1.3%). Beyond basic cell type identification, the knowledge-based approach revealed multi-scale tissue organization including 27 cell state transitions (such as granulosa-to-theca transitions with 3,306 cells showing intermediate profiles), 4 significant pathway co-expression networks (steroidogenesis *↔* folliculogenesis: 0.433, folliculogenesis *↔* cell cycle: 0.483, folliculogenesis *↔* hormone signaling: 0.352, ovulation hormone signaling: 0.533), and 13 spatially coherent functional domains where specific biological processes operate at high intensity (3 steroidogenesis domains with 2,410 cells, 7 folliculogenesis domains with 2,874 cells, 3 cell cycle domains with 746 cells). This shows that integrating biological knowledge captures not only cell type identity but also functional states, developmental trajectories, and spatial functional architecture.

AI-Assisted Biological System generated and validated 3 biological hypotheses with 100% success rate, achieving 3 high-confidence discoveries in a single iteration. The agent employed systematic pattern detection to generate one foundational hypothesis and GPT-4o-mini to generate two creative hypotheses connecting observed patterns to biological mechanisms: microenvironmental influence on expression despite absence of spatial clustering, and physiological regulation explaining sparsity patterns. GPT-4o-mini provided detailed biological interpretations contextualizing computational metrics within biological knowledge—for example, explaining that moderate silhouette score (0.4) indicates biologically meaningful clustering despite some overlap, and connecting sparse expression to spatiotemporal gene regulation rather than technical limitations. The system demonstrated intelligent decision-making through early termination, recognizing sufficient evidence after one iteration rather than exhausting all planned iterations. This work shows that large language models can effectively augment quantitative spatial transcriptomics analysis with creative hypothesis generation and biological interpretation, reducing manual effort while enhancing discovery through AI-driven reasoning that connects computational findings to mechanistic biological understanding.

### 6.2 Practical Application for Researchers

This framework is directly applicable to translational research settings where distinguishing spatially co-located but biologically distinct cell populations is critical. For example, in tumor biopsy analysis, a researcher could apply the knowledge-based annotation pipeline to a spatial transcriptomics dataset of mixed tumor tissue and receive spatially-resolved maps that separate malignant epithelial cells from benign stromal or immune cells within the same tissue section. The cell state transition detection would further highlight boundary zones where benign cells show intermediate expression profiles consistent with early malignant transformation, providing spatially actionable targets for biopsy or therapeutic intervention. The pathway co-expression analysis could then identify which signaling hubs are activated specifically in the malignant regions, helping prioritize drug targets with maximal spatial specificity.

### 6.3 Future Work

#### Ensemble Clustering Methods

Each of current clustering algorithms chosen in this project represents the most basic form within its respective family. Future work could investigate more advanced alternatives within each family to better capture the complexity of spatial transcriptomics data. Within the partitioning-based family, K-means could be replaced or supplemented by Partitioning Around Medoids (PAM), which is more robust to outliers since it uses actual data points as cluster centers rather than means. Within the hierarchical family, divisive methods such as DIANA could be explored alongside the current agglomerative approach, as top-down splitting may better reflect the biological hierarchy of cell type organization in tissue. Within the model-based family, the standard Gaussian assumption could be relaxed by adopting heavier-tailed distributions such as the multivariate *t*-distribution, contaminated normal distributions, or the Generalized Hyperbolic Distribution (GHD), which are better suited to handle the outliers and skewness commonly observed in gene expression data. More broadly, the choice of clustering algorithm should be guided by the characteristics of the dataset, as different tissue types, sequencing platforms, and sparsity levels may favour different algorithmic families. Future work could therefore develop a data-adaptive algorithm selection strategy that matches the most appropriate clustering methods to the specific attributes of each spatial transcriptomics dataset before constructing the ensemble.

#### Knowledge-Based Annotation

For knowledge-based clustering, expanding annotation by curating additional markers to improve coverage rate and integrating multi-omics data (spatial proteomics, metabolomics) would allow experimental validation of the 27 computationally predicted cell state transitions and confirm pathway activities beyond transcriptional inference alone.

#### AI Agent

For the autonomous agent, learning from multiple datasets to develop analysis strategies that generalise across different tissues and disease contexts can potentially improve overall analytical performance and hypothesis quality.

#### Biomedical and Drug Research Applications

The framework has significant potential for translational biomedical research. In disease research, the spatial cell type annotation and pathway activity mapping could be applied to tumour microenvironment analysis, enabling identification of cancer-associated cell states and spatially organised immune infiltration patterns relevant to immunotherapy response. The cell state transition detection could help characterise disease progression trajectories, for example identifying intermediate cell states in fibrosis or neurodegeneration that represent actionable therapeutic windows. In drug discovery and development, the pathway co-expression analysis could be used to identify candidate drug targets by pinpointing spatially localised signalling hubs where intervention would have maximal effect while minimising off-target impact. The autonomous hypothesis generation agent could accelerate target identification by rapidly screening large spatial transcriptomics datasets from patient biopsies and generating ranked, testable hypotheses about mechanism of action. As spatial transcriptomics becomes more widely adopted in clinical trial tissue profiling, this framework could support biomarker discovery by characterising how spatial tissue organisation correlates with treatment response, providing a more comprehensive picture than bulk or single-cell approaches alone.

## Availability

The complete implementation is publicly available at https://github.com/wandreopoulos/agentic_spatial_transcriptomics_analysis. Source code, documentation, and example configurations are openly available under the MIT license.

## Acknowledgments

Computational resources were provided by the SJSU computing infrastructure.

## Contributions

This project was completed as part of a Master of Science, Data Science, by Maiqi (Maggie) Zhang in the Department of Computer Science at San José State University. MZ developed the code, discovered datasets, performed experiments, interpreted the results, wrote the manuscript. Professor WBA provided mentorship, ideas, and feedback throughout the project and edited the final version. Professor CP provided comments, feedback and committee guidance. MR developed and maintains the cleaned-up version of the software.

